# Virtual Liver Needle Biopsy from Reconstructed Three-Dimensional Histopathological Images: Quantification of Sampling Error

**DOI:** 10.1101/2022.06.08.495226

**Authors:** Qiang Li, Fusheng Wang, Yaobing Chen, Hao Chen, Shengdi Wu, Alton B. Farris, Yi Jiang, Jun Kong

## Abstract

**Introduction:** Prevalently considered as the “gold-standard” for diagnosis of hepatic fibrosis and cirrhosis, the clinical liver needle biopsy is known to be subject to inadequate sampling and a high mis-sampling rate. However, quantifying such sampling bias has been difficult as generating a large number of needle biopsies from the same living patient is practically infeasible. We construct a three-dimension (3D) virtual liver tissue volume by spatially registered high resolution Whole Slide Images (WSIs) of serial liver tissue sections with a novel dynamic registration method. We further develop a Virtual Needle Biopsy Sampling (VNBS) method that mimics the needle biopsy sampling process. We apply the VNBS method to the reconstructed digital liver volume at different tissue locations and angles. Additionally, we quantify Collagen Proportionate Area (CPA) in all resulting virtual needle biopsies in 2D and 3D.

**Results:** The staging score of the center 2D longitudinal image plane from each 3D biopsy is used as the biopsy staging score, and the highest staging score of all sampled needle biopsies is the diagnostic staging score. The Mean Absolute Difference (MAD) in reference to the Scheuer and Ishak diagnostic staging scores are 0.22 and 1.00, respectively. The absolute Scheuer staging score difference in 22.22% of sampled biopsies is 1. By the Ishak staging method, 55.56% and 22.22% of sampled biopsies present score difference 1 and 2, respectively. There are 4 (Scheuer) and 6 (Ishak) out of 18 3D virtual needle biopsies with intra-needle variations. Additionally, we find a positive correlation between CPA and fibrosis stages by Scheuer but not Ishak method. Overall, CPA measures suffer large intra- and inter-needle variations.

**Conclusions:** The developed virtual liver needle biopsy sampling pipeline provides a computational avenue for investigating needle biopsy sampling bias with 3D virtual tissue volumes. This method can be applied to other tissue-based disease diagnoses where the needle biopsy sampling bias substantially affects the diagnostic results.

## 1 Introduction

Chronic liver disease has a high and steadily increasing mortality rate worldwide [1]. As the disease progresses, the abnormal deposition of excess extracellular matrix (ECM) rich in fibrilforming collagens leads to liver fibrosis, an important pathology hallmark in a variety of liver diseases, especially for chronic liver disease [2]. The accumulation of the ECM over time can result in distortion of hepatic architecture and eventually in cirrhosis that is accompanied by stiffening of liver tissues, loss of liver functions, hepatocellular carcinoma, and portal hypertension [2].

In current clinical practice, liver needle biopsy is the gold standard for diagnosis and therapeutic planning [3]. Typical core biopsies are 10 − 30*mm* by 1 − 2*mm*, representing only about 0.002% of the total liver mass [4]. Therefore, the needle biopsy based diagnostic results highly depend on such factors as the sampling location, direction, depth, and number. However, there is a lack of practical protocols to optimize these sampling parameters in a clinical setting [5, 6, 7]. Additionally, such biopsies for diagnosis often produce very few two-dimensional (2D) slides for pathologists to review under a microscope [8, 9, 10]. The histology information captured in such 2D needle biopsy slides, therefore, is subject to a large sampling bias, contributing to erroneous disease classification and staging results in the downstream analyses. Compared to the whole liver disease representation, small-scaled needle biopsies are inadequate in capturing tissue alterations, which is the major source of the sampling error [11].

As pathologists make their diagnoses based on the biopsies, it is a critical to assess the biopsy sampling bias and its impact on the resulting diagnostic results. However, there is a lack of feasible methods to carry out such sampling bias studies, as it is not feasible to take numerous needle biopsies from the same patient just to support such research. Compared to the limited biopsy samples for diagnosis, resected tissues after surgeries can provide much more histopathological information at a much larger volume ideal for the sampling bias study. Although clinical experiments have reported a high rate of sampling bias in needle liver biopsies [11, 12], no generic, fully automated, or computational tools exist to enable quantitative biopsy sampling heterogeneity investigations. To address this critical problem, we have investigated the needle biopsy sampling bias by digitally generating multiple virtual needle biopsies from a digital three-dimension (3D) liver tissue volume. In this study, adjacent 2D tissue slides of a large resected 3D tissue structure are stained and scanned by a high throughput digital scanner to generate serial histopathology Whole Slide Images (WSIs). Such serial WSIs can collectively encode lossless histology information. Using these imaging data, we explore the liver needle biopsy sampling variation by generating numerous virtual liver needle biopsies at multiple tissue locations and directions.

Recent advances in tissue engineering have introduced novel 3D tissue volumes [13]. Thanks to the advance of high throughput digital scanners, histopathology WSIs of liver tissue biopsies with the Masson’s trichrome stain can be efficiently produced on a daily basis. With such 2D imaging data, unfortunately, the structural and quantity information of liver collagen can differ from the 3D space. To extend the 2D histopathology analysis to the information-lossless 3D space, image registration methods are required [14, 15, 16, 17, 18]. However, current methods for pathology image registration lack the capability for processing WSIs at the highest image resolution and may result in computer memory overflow issue as a single WSI can have giga-pixels of data [19, 20, 21, 22, 23, 24, 25, 26, 27]. Overall, generation of such 3D bright-field WSI tissue section volumes from serial 2D WSIs remains a significant technical challenge as each WSI can have a giga-pixel image resolution.

Additionally, manual histopathology component analyses are often extremely time-consuming due to the overwhelmingly large image scale. By contrast, computer-based image processing techniques are much more scalable and subject to less inter- and intra-reader variability. Therefore, there is a growing demand to use image processing methods to efficiently detect and count image pixels representing liver histology components in the entire biopsy images for an enhanced processing throughput, reliability and reproducibility [28, 29, 30, 31, 32].

In this study, we develop a novel computational analysis pipeline to assess the liver needle biopsy sampling error. Illustrated in Figure 1, the developed system The method consists of 1) 3D digital liver tissue volume reconstruction with spatially aligned WSIs of serial tissue resections, 2) Virtual Needle Biopsy Sampling (VNBS) function mimicking the physical needle biopsy sampling process in the clinical practice, 3) automatic quantification of liver Collagen Proportionate Area (CPA) from such 3D virtual needle biopsies, and 4) quantitative inter- and intra-needle biopsy sampling error analysis. Additionally, our 3D liver CPA quantification analysis demonstrates a promising potential to aid pathologists for diagnostics in an authentic 3D tissue space. Overall, our method provides a new and generic computational tool to support biopsy sampling investigations critical to the biopsy based diagnostic reviews of a large spectrum of diseases.

**Figure 1:**
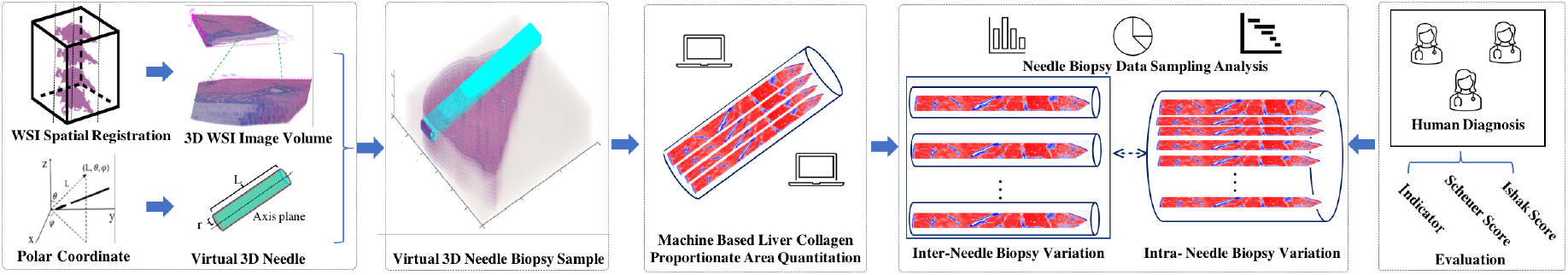
The overall framework of the developed 3D virtual liver needle biopsy sampling quantification pipeline. A 3D virtual liver tissue volume is first constructed by spatially registered high resolution Whole Slide Images (WSIs) of serial liver tissue sections with a dynamic registration method. The developed Virtual Needle Biopsy Sampling (VNBS) function mimicking the clinical needle biopsy sampling process is applied to the reconstructed digital liver image volume for multiple times, each at a distinct tissue location and angle. All 3D virtual needle biopsy samples are graded by a board-certified pathologist using the Scheuer and Ishak staging systems. Additionally, Collagen Proportionate Area (CPA) in all resulting virtual needle biopsies in 2D and 3D are quantitated. Finally, this pipeline provides an analysis module for quantitative inter- and intra-needle biopsy sampling variation investigations.

## 2 Methods and data

### 2.1 Liver tissue preparation and imaging

Human liver tissues were prepared and scanned with the standard tissue preparation protocols, at Wuhan Tongji Hospital, China. The liver tissues were cut into ten 0.5 *×* 0.5 *cm*^2^ serial sections, embedded into paraffin blocks, and stained with Masson’s trichrome stain for histological assessment. The blocks were sectioned into slides at 4 µm thickness. The tissue was mordanted in Bouin’s solution for 15 minutes. The slides were washed in running tap water to remove all the picric acid from sections. The sections were stained in Weigert’s working hematoxylin for 10 minutes. After sections were washed again, they were stained in Biebrich scarlet acid fuchsin. The slides were stained in Phosphotungstic/Phosphotungstic acid solution and rinsed in distilled water three times before placed in 1% acetic acid solution for one minute. The slide images were scanned using a 40X objective lens (NanoZoomer S360, Hamamatsu Photonics, Hamamatsu, Japan). The physical resolution of the resulting WSIs is 4.437 *×* 10^−1^ µm per pixel. Each image was acquired from a full-size liver tissue in the slide, where normal hepatic cells, collagen fibers and blood vessels are highlighted by pink, blue, and white color.

### 2.2 Registration of full resolution WSIs of serial liver tissue sections

We have used a dynamic multi-resolution registration method to construct a 3D tissue from 2D serial slides [33]. As demonstrated in Figure 2, we first align serial tissue structures at a low resolution with a landmark-based rigid transformation [34] and a non-rigid registration to simultaneously maximize the normalized mutual information and minimize the transformation energy [16, 35]. The middle WSI in the serial slide sequence serves as the reference image that all other WSIs are mapped to. The reference image is partitioned into non-overlapping image patches of a smaller size such that the computer memory can readily accommodate each image patch individually and dynamically. Next, we map the rigid and non-rigid transformations derived from the low resolution images to the full resolution WSIs. For serial WSI registration at the full image resolution, an iterative transformation propagation method is used to align nonreference images to the common reference image [33].

**Figure 2:**
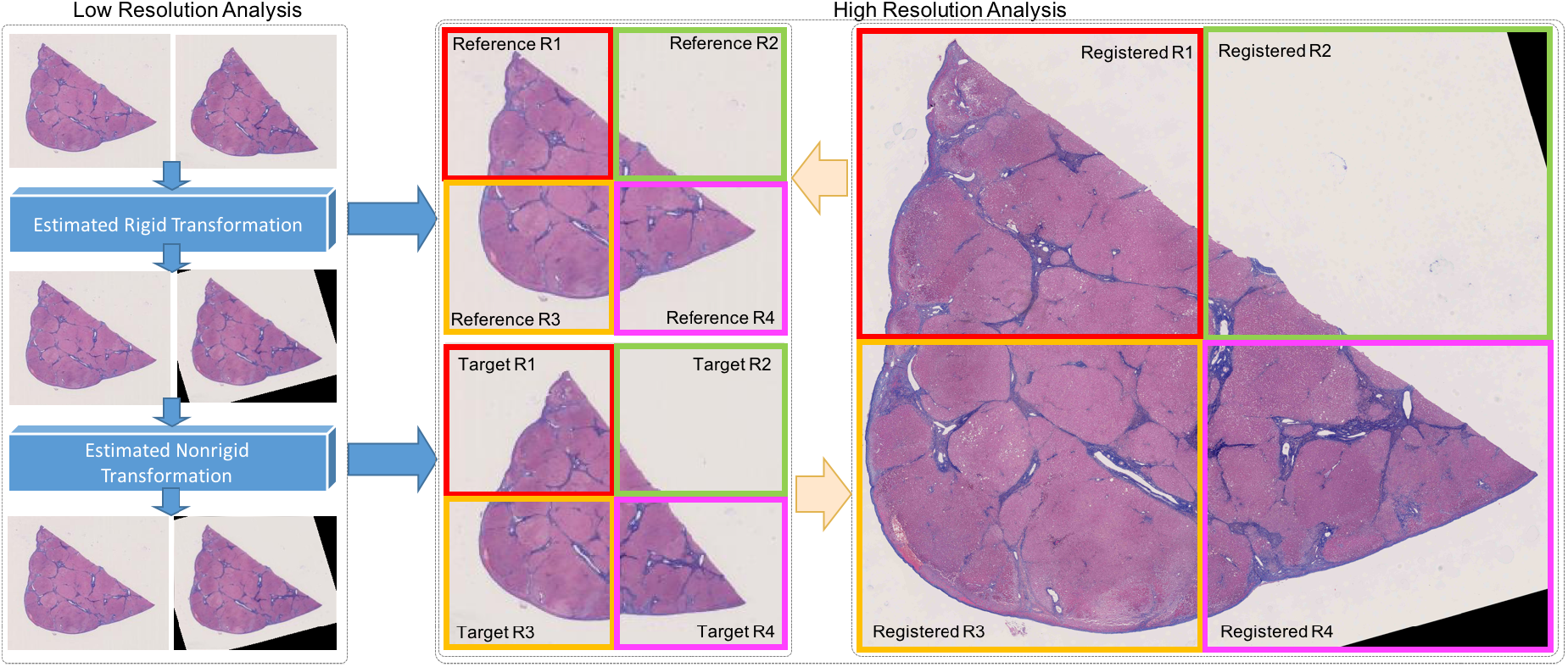
The schema of the employed registration method processing full resolution WSIs of serial liver tissue sections. Registration of serial WSIs is completed by two steps: estimation of rigid and nonrigid transformations of paired WSIs at a low image resolution; and image patchwise dynamic registration at the full WSI resolution with mapped and aggregated high resolution spatial transformations derived from low image resolution spatial transformations. At the full WSI resolution, the reference WSI is partitioned into image patches, making it feasible for computer memory accommodation and the efficient parallel computation.

### 2.3 Virtual 3D needle biopsy sampling

Most common liver biopsy needles in current clinical practice are either for diffuse disease investigations with a core diameter greater than 1.0 mm or for focal lesion investigations with the internal diameter less than 1.0*mm* [36]. In this study, we digitally simulate the small needle biopsy using a 16-gauge biopsy needle. The internal radius of the 16-gauge needle is about 700 µm, with a length of about 20*mm* [37]. Each 3D virtual needle biopsy has 45,000 voxels in length (20*mm*) and 1,600 voxels in radius (700 µm), as each image pixel takes 0.4437 µm by each dimension. To implement the 3D virtual needle biopsy sampling, we develop a Virtual Needle Biopsy Sampling (VNBS) function. As all points in a 3D virtual needle biopsy of a given specification can be readily represented by the spherical coordinate system, the process for a 3D virtual biopsy needle penetrating a 3D tissue volume can be characterized by spatial transformations. A 3D virtual needle biopsy can be described by the VNBS function by a set of parameters: (*P*_0_, *L, R, θ, φ*). *P*_0_ is the center of the needle tip cross-section at the tissue entry. The needle length and the radius of the cross section area are represented by L and R, respectively. In the spherical coordinate system (*L, θ, φ*) in Figure 3(a), we represent a 3D virtual biopsy needle sampling process by a spatial transformation:

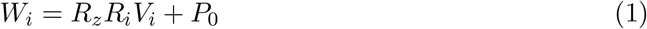

where *V*_*i*_ is the coordinate mesh-grid of a virtual 3D needle and two rotation matrices are given as:

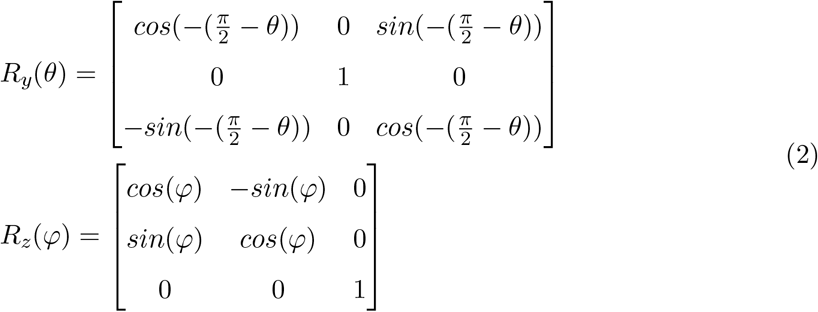

The resulting 3D needle biopsies are illustrated in Figure 3(b). Two sets of experiments are performed to validate the VNBS function for 3D tissue volumes. First, we produce a synthetic data volume, with 300 × 300 × 600 voxels and a total of 12 cuboids in distinct colors. An array of 3D virtual needle biopsy samples over this synthetic data volume is presented in Figure S1, with the resulting 2D longitudinal image plane from each 3D needle biopsy. Second, we apply the VNBS function to a 1, 000 × 1, 000 × 50 semi-synthetic image volume with different needle locations and angles. The resulting 3D needle biopsies and their 2D longitudinal planes are illustrated in Figure S2. After careful visual examinations of the results from these two experiments, we conclude the 3D virtual needle biopsies match well with the ground truth.

**Figure 3:**
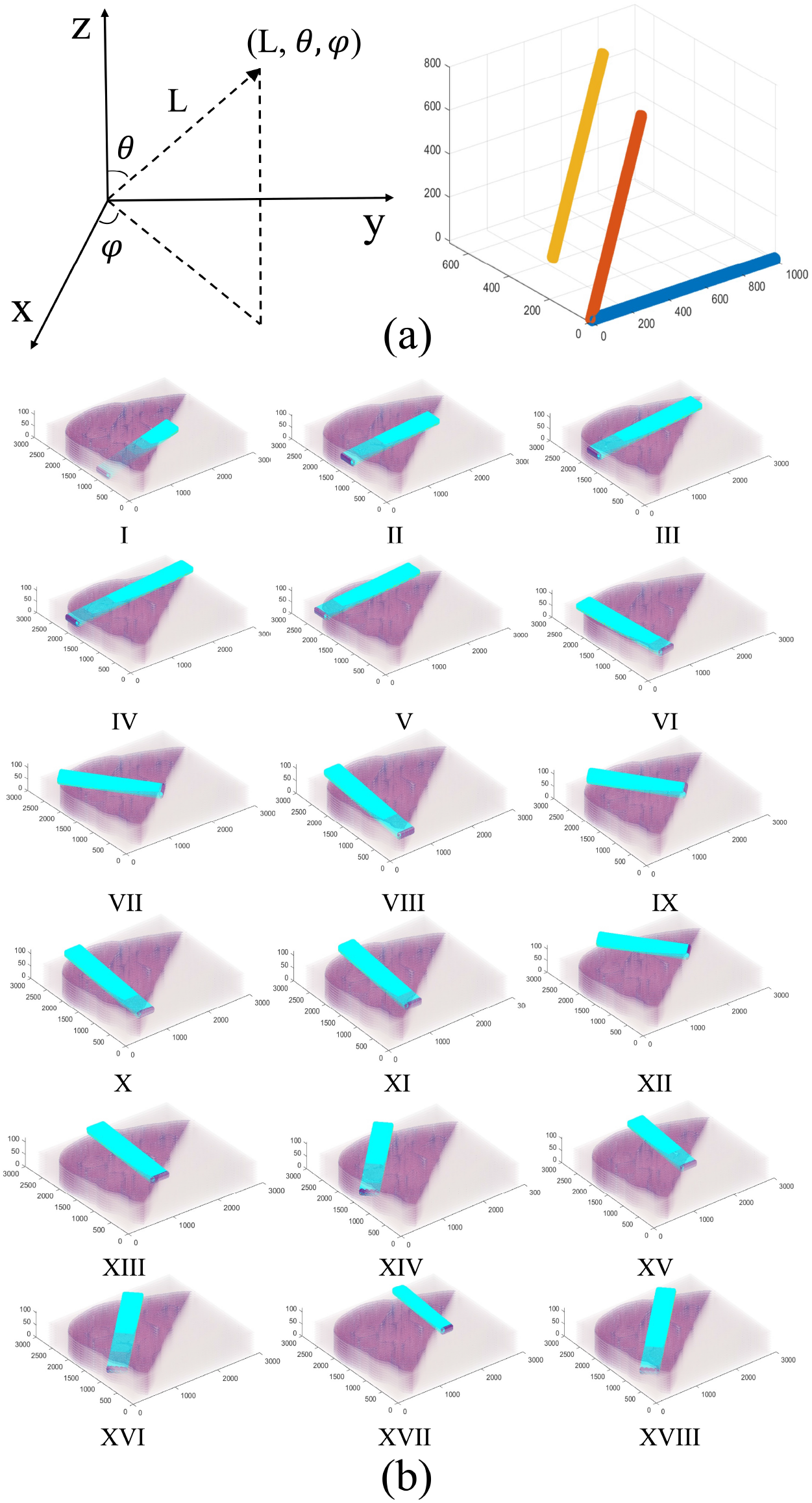
Virtual 3D needle biopsy sampling process. (a) Virtual 3D needle parameters are presented in a spherical coordinate system where the initial, rotated and translated biopsies are illustrated in blue, orange, and yellow, respectively; (b) Randomly produced 3D needle biopsies are visually plotted after they are down scaled.

### 2.4 Generation of 2D longitudinal image planes from virtual 3D needle biopsies

In practice, pathologists review 2D longitudinal biopsy cross-sections that are often arbitrarily chosen. To determine if cross-sectioning of the 3D biopsy sample influences the tissue grading by pathologists, we generate 11 longitudinal cross-sections from each 3D virtual needle biopsy. We apply a 3D linear interpolation to create 2D longitudinal planes within a 3D virtual needle biopsy [38]. We rotate the 2D longitudinal image planes in a 3D virtual needle biopsy to generate multiple cross-section images rotated around the longitudinal needle axis. We denote *P* = *{*(*x, y, z*)|0 ≤ *x* ≤ *L*, −*R* ≤ *y* ≤ *R, z* = 0*}*as a set of voxel centers on the x-y plane. Discrete voxel centers in the longitudinal plane by a rotation angle *α* around the longitudinal needle axis can be found by:

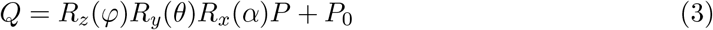

where

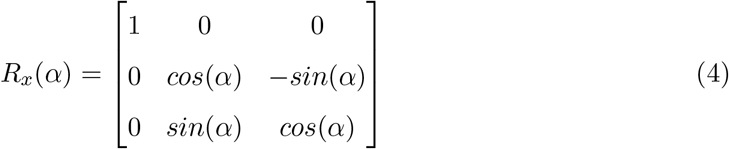

The longitudinal cross-sectional plane is parallel to the x-y plane when *α* = 0, and perpendicular to the x-y plane when *α* = 90^°^. Varying *α* allows us to generate different longitudinal planes from the same 3D virtual needle biopsy and visualize the internal tissue details of these longitudinal cross sections. Supported by this method, the intra-biopsy variance is studied with image planes of diverse rotation angles along the longitudinal needle axis. We confirm the quality of the 2D cross-sections from 3D virtual needle biopsies using the same semi-synthetic tissue volume at *θ* = 90^°^, *φ* = 0^°^, or 90^°^ as we know what the resulting 2D longitudinal image planes should look like by these rotation angles. Additionally, we compare different interpolation methods, including ‘linear’, ‘nearest’, ‘cubic’, ‘spline’, and ‘makima’ (i.e. Modified Akima cubic Hermite interpolation), and choose the linear interpolation for its good interpolation quality and small execution time cost. We present interpolated 2D image planes rotated around the 3D virtual needle axis by *α* = 0^°^, 30^°^, 60^°^, and 90^°^ in Figure S3, respectively.

### 2.5 Liver collagen quantification based on Masson trichrome stain

As liver disease staging and prognostics depend on liver collagen assessments [39], we have developed a complete 3D liver collagen quantification pipeline with multiple processing modules (Figure 4). For the registered WSIs stained with Masson trichrome, we first normalize the images to reduce the impact of the stain variation on collagen quantification. Next, we decompose the color channels by the color deconvolution and focus on collagen signal channel for further collagen quantification. Figure 4(a) illustrates the two deconvolved color channels and the original image region. The resulting color channels are converted to grayscale images that are enhanced by intensity contrast [40], adaptively thresholded for binary image production [31] and finally represented in pseudo-color in Figure 4(b). Figure 4(c) illustrates the binary images of the deconvolved blue and purple stain signals, and pseudo-colored label images of tested serial WSIs. CPA is defined as the ratio between the number of voxels with the liver collagen label (i.e. blue in pseudo-color) and that of voxels labeled as other tissue component (i.e. red in pseudocolor) in a 3D virtual needle biopsy sample.

**Figure 4:**
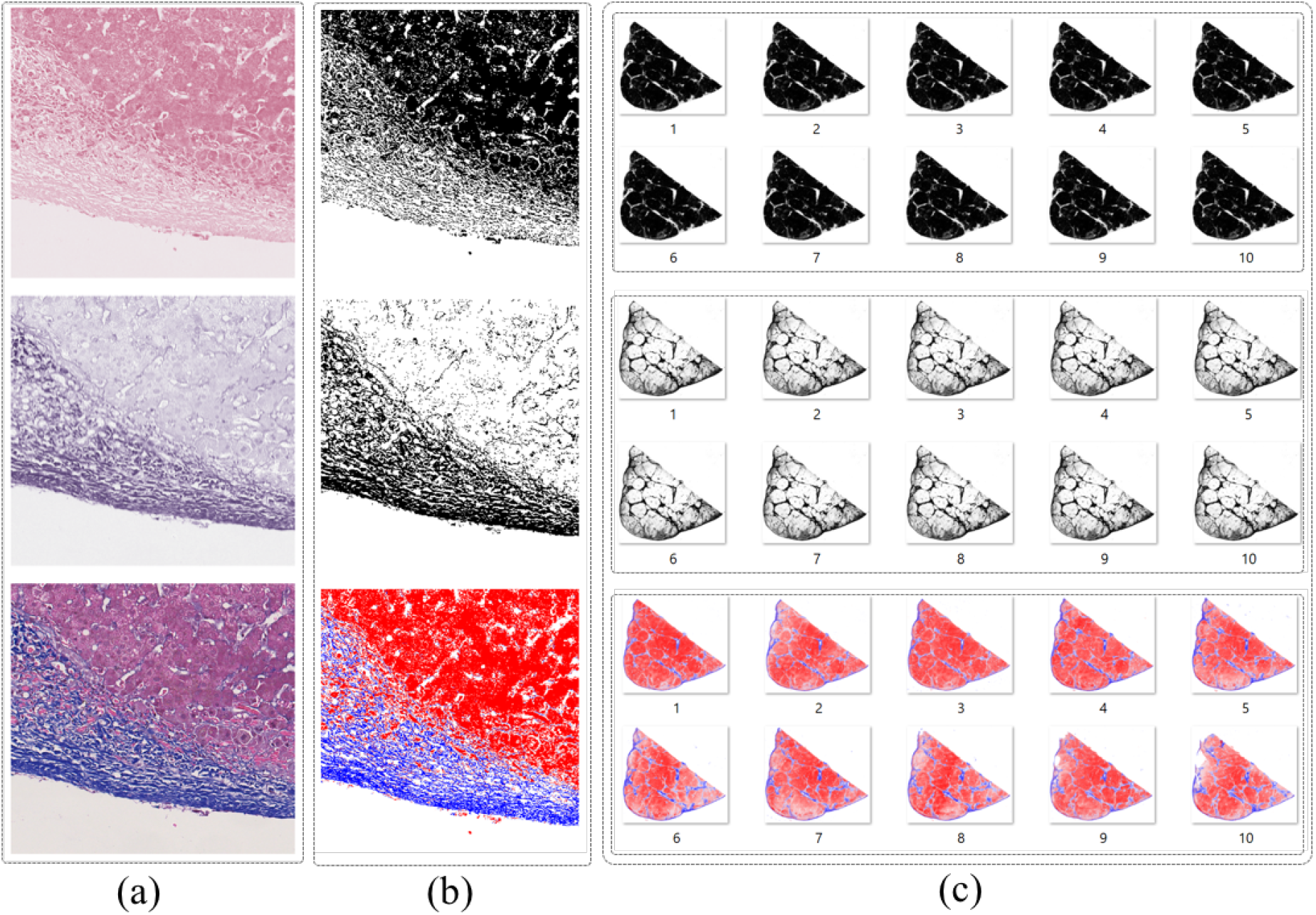
Collagen quantification by the color deconvolution with serial WSIs of liver tissues in Masson’s trichrome stain. (a) We present the deconvolved color channel for noncollagen component in pink (top), liver collagen in blue and purple (middle), and the original image with mixed signals (bottom); (b) The decomposed color channels in (a) are converted to binary images (top and middle) that are jointly represented by a label image in pseudo-colors, i.e. blue for collagen and red for other tissue components (bottom); (c) Binary images for two stain signals and pseudo-colored label images of serial WSIs in our dataset are presented from top to bottom.

### 2.6 Liver fibrosis staging

All biopsy image planes from random 3D virtual liver needle biopsies are evaluated by an experienced pathologist with Scheuer and Ishak scoring systems [28, 41, 42]. With this reviewing process, each 2D cross-sectional image of the 3D virtual biopsy receives two staging scores, the 5-tier Scheuer score ranging between 0 and 4, and the 7-tier Ishak score ranging between 0 and 6.

## 3 Results

### 3.1 Virtual 3D liver tissue volume reconstruction with serial WSIs

By the WSI registration method, all WSIs of serial liver tissue sections in our dataset are registered to the common reference image space in the full resolution. we crop the non-tissue part to minimize the volume of non-interest. Demonstrated in Figure 5(a), the resulting image resolution of each registered WSI is 28, 832 *×* 26, 400 pixels. Additionally, 3D rendering of these raw, rigidly registered, and non-rigidly registered WSIs are illustrated in Figure 5(b), with a close-up view of a small image volume shown on the right panel. As these serial tissue sections are physically cut during the tissue preparation process, the slice thickness is larger than the x-y in-plane pixel size. Therefore, the inter slice interpolation is used to generate the intermediate slides along the z direction. The resulting 3D liver tissue volume has 28, 832 × 26, 400 × 110 voxels.

**Figure 5:**
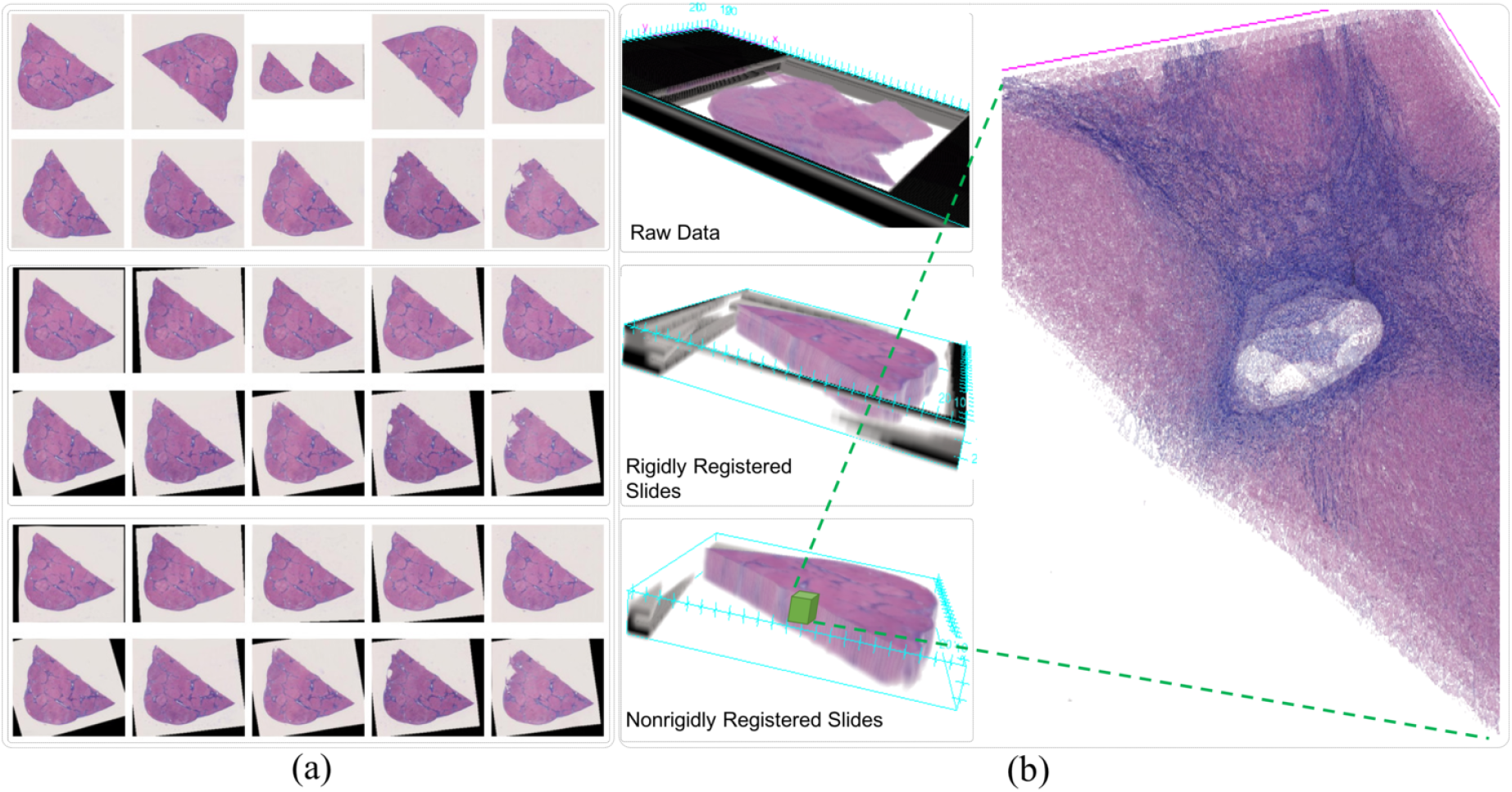
Construction of a 3D liver tissue volume from serial WSIs by image registration. (a) Panels of ten original, rigidly registered, and nonrigidly registered serial WSIs of liver tissue sections are presented from top to bottom; (b) 3D volume rendering visualizations with original, rigidly registered, and nonrigidly registered serial WSIs of liver tissue sections are illustrated from top to bottom. Additionally, the close-up view of a small registered tissue volume from the nonrigidly registered image volume is presented on the right. After the background image volume is trimmed, the image resolution of each registered slide is 28, 832 *×* 26, 400 by pixels. After data interpolation in the resulting 3D tissue space, the final 3D image volume has 28, 832 *×* 26, 400 *×* 110 voxels.

### 3.2 CPA quantification and collagen visualization from a 3D liver tissue volume

With the constructed 3D live tissue volume, two stain signal channels are deconvolved and presented in Figure 6(a). The resulting pseudo-colored label images for liver collagen fibers and the reconstructed 3D liver collagen volume are illustrated in Figure 6(b). The overall CPA from the whole 3D liver tissue volume is 14.5%. Such 3D liver collagen volumes can provide a more straightforward visual understanding of the liver collagen spatial distributions in a 3D tissue space. By our analysis results, we also notice the fibers show marked bridging (portal-portal and/or portal-central) with nodules, consistent with the definition of stage 3 by Scheuer and 5 by Ishak staging system, respectively.

**Figure 6:**
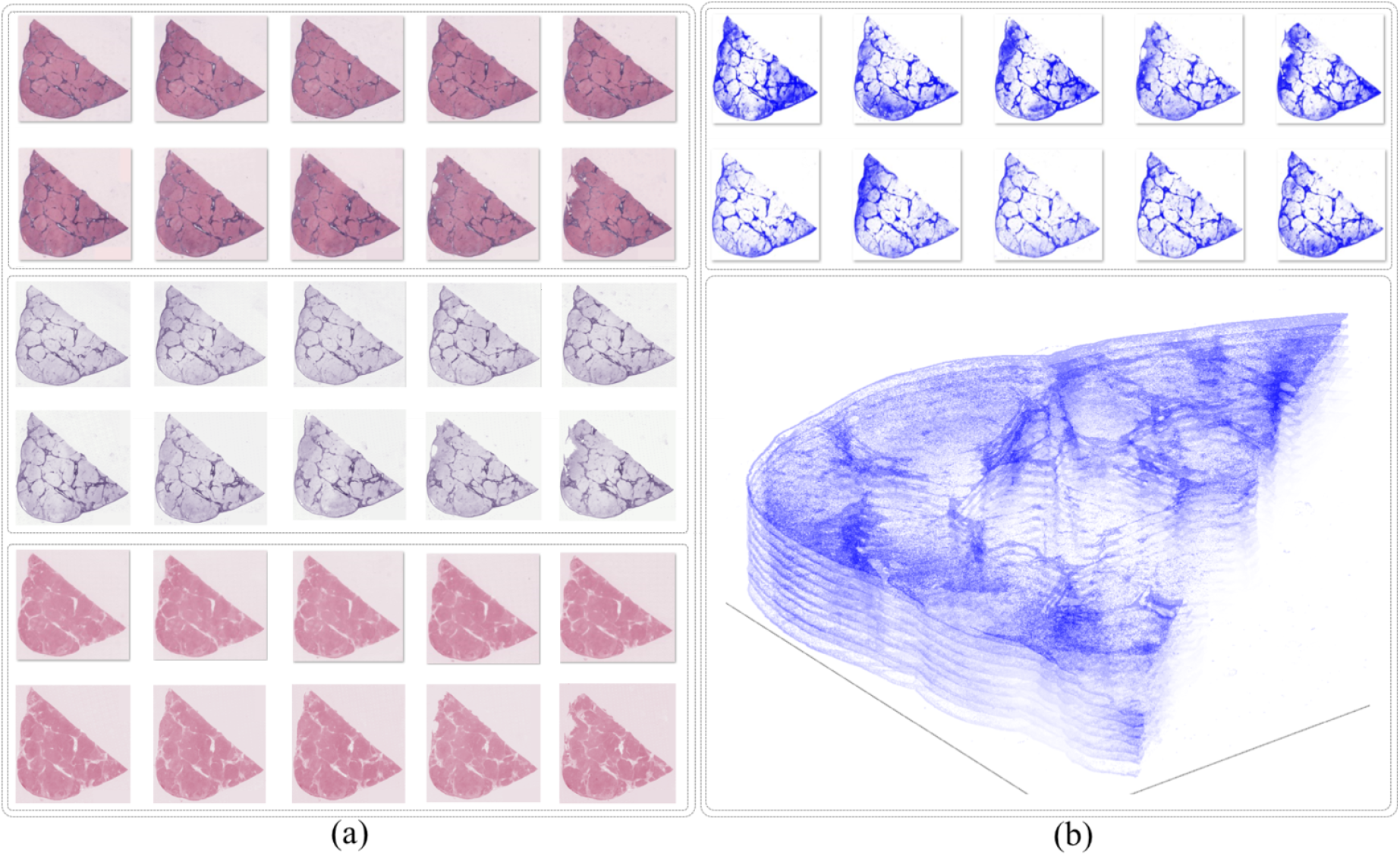
Construction of a 3D liver collagen volume from serial WSIs by color deconvolution. (a) The normalized serial WSI set, and its decomposed two stain signal sets are presented from top to bottom; (b) The resulting pseudo-colored label images for liver collagen and the constructed 3D liver collagen volume are illustrated.

### 3.3 Virtual 3D needle biopsy generation

Although we can generate as many virtual needle biopsies as possible from the 3D tissue volume by the VNBS function, we use n=18 virtual needle biopsies due to their sufficient tissue representation power. To determine the optimal number of biopsies, we vary the sampling number in simulations. For each fixed number of biopsies n, we repeat the virtual sampling process, the resulting needle biopsy CPAs are compared with the overall tissue volume CPA by the t-test. We use the percentage of the sampling processes with the p-value ¡ 0.05 as a measure of tissue heterogeneity representation power. Such measure is plotted as a function of the needle biopsy number in Figure S4. It is noticed that the tissue representation measure saturates rapidly (¿95%) as the biopsy number increases to 18, suggesting a sufficient tissue representation power. The resulting liver virtual biopsies are illustrated in Figure 7(a) by ascending CPA measures.

**Figure 7:**
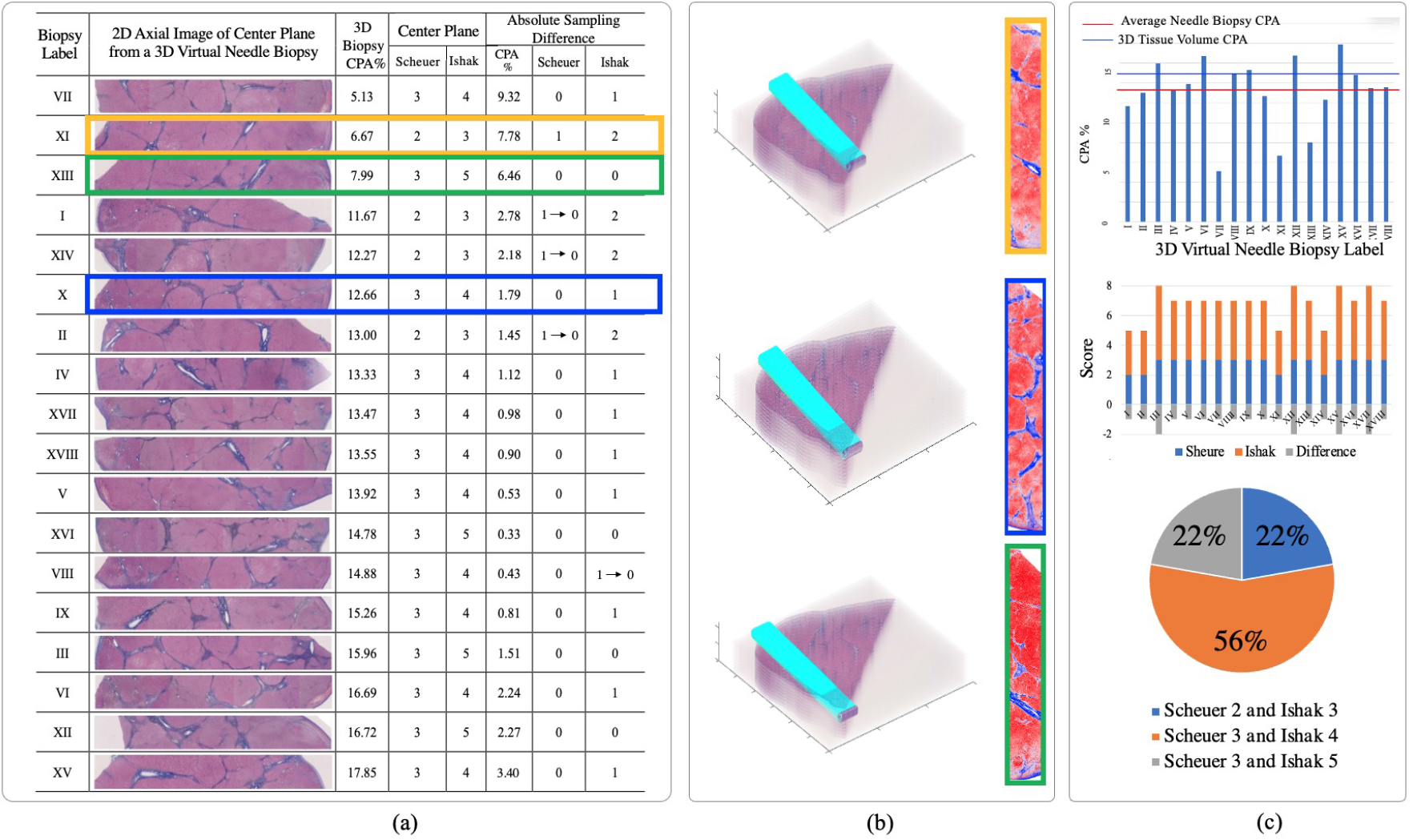
Liver staging and Collagen Proportionate Area (CPA) quantification with virtual 3D needle liver biopsies. The developed Virtual Needle Biopsy Sampling (VNBS) function is used to randomly sample virtual 3D needle biopsies at distinct positions and angles. The resulting 3D biopsies are manually graded by Scheuer and Ishak histological staging method, and automatically quantitated for liver CPA. 2D longitudinal images from 3D virtual needle biopsies, machine based CPA quantitation, and human grading results on 2D longitudinal images are demonstrated in (a) by the CPA ascending order, along with the sampled liver biopsy CPA difference from the overall 3D tissue volume CPA, and absolute sampling biopsy staging score differences. Mimicking clinical practice, we use the staging score of the center 2D longitudinal image plane as the biopsy staging score. Additionally, we take the highest staging score of all sampled needle biopsies as the diagnostic staging score. Representative 3D biopsies of distinct combinations of Scheuer and Ishak staging scores, i.e. (2, 3), (3, 4), and (3, 5), are demonstrated in (b), along with the segmented axial image planes with collagen in blue and other tissue components in red, respectively. The Machine based CPA results of sampled virtual needle biopsies, their Scheuer and Ishak staging scores, and the percentage of 3D needle biopsies of distinct combinations of Scheuer and Ishak staging scores are presented in (c).Interestingly, when we define the biopsy staging score as the highest staging score of all 2D longitudinal image planes within a 3D virtual needle biopsy, the numbers of sampling needle biopsies with biopsy staging scores different from the diagnostic score are reduced by both staging methods, as suggested by the arrow in the right most two columns in (a).

### 3.4 Inter-3D virtual needle biopsy sampling heterogeneity

#### Inter-3D virtual needle biopsy heterogeneity by staging scores

By the Scheuer and Ishak histological staging methods, all cross-section images of all 3D virtual needle biopsies are graded by an experienced pathologist. Following the clinical reviewing process, we take the staging score of the center 2D longitudinal image plane with the rotation angle 0^°^ as the biopsy staging score. Additionally, we take the highest staging score of all sampled needle biopsies as the diagnostic staging score. Characterized in Figure 7(a), the sampling difference is evaluated by the absolute differences between individual biopsy staging scores and the diagnostic staging score. The Mean Absolute Difference (MAD) in reference to the Scheuer and Ishak diagnostic staging scores are 0.22 and 1.00, respectively. Additionally, 22.22% and 77.78% of virtual biopsies show lower biopsy staging scores than the diagnostic score by the Scheuer and Ishak staging methods, respectively. Specifically, the absolute Scheuer staging score difference in 22.22% of sampled biopsies is 1. By the Ishak staging method, 55.56% and 22.22% of sampled biopsies present score difference 1 and 2, respectively. Interestingly, when we redefine the biopsy staging score as the highest staging score of all 2D longitudinal image planes within a 3D virtual needle biopsy, the MAD is decreased to 0.06 and 0.94. Additionally, sampling needle biopsies with biopsy staging scores different from the diagnostic score are reduced to 5.56% and 72.22% of all 3D virtual needle biopsies by Scheuer and Ishak staging methods, respectively.

Additionally, different combinations of Scheuer and Ishak staging scores are observed in sampled virtual needle biopsies. Figure 7(b) demonstrates representative 3D virtual needle biopsies (i.e. virtual biopsy X, XI, and XIII) with slightly shifted tissue sampling entry points. The staging score pairs by Scheuer and Ishak method for these three needle biopsies are (3, 4), (2, 3) and (3, 5), respectively. Therefore, such small spatial changes in the tissue entry point lead to distinct score combinations. By contrast, virtual 3D needle biopsies with diverse sampling locations and angles from the same 3D tissue imaging volume can share the same staging score combinations by Scheuer and Ishak method. For example, virtual needle biopsy sets (I, XI, XIV), (IV, VII, X), (III, XIII,XVI) present Scheuer-Ishak staging score combinations of (2, 3), (3, 4), and (3, 5), respectively. In each such set, the sampling locations and angles are significantly different. These results suggest that the sampling difference can substantially affect the staging score.

#### Inter-3D virtual needle biopsy heterogeneity by CPAs

CPAs from 3D liver needle biopsies and their absolute difference from the overall 3D tissue volume derived CPA are presented in Figure 7(a). The overall liver tissue CPA from the reconstructed virtual 3D tissue volume is 14.5%, while the average CPA from our 18 virtual 3D liver needle biopsies is 13.1% with a standard deviation 0.8%. The detected collagen fibers and other tissue components in the 2D longitudinal image planes from these representative virtual needle biopsies are represented in blue and red in Figure 7(b). Additionally, we present needle biopsy CPA quantitation results in Figure 7(c). The resulting individual needle biopsy CPAs present a relatively large variation in reference to the average biopsy CPA (i.e. 13.1% by the red line), with the resulting MAD reaching 27% of the average biopsy CPA.

#### Inter-virtual needle biopsy heterogeneity using 2D longitudinal biopsy images

From each 3D virtual needle biopsy, we generate 11 2D longitudinal image planes with the rotation angle 0°, ±0.5°, ±1°, ±1.5°, ±2°, and ±2.5°. A set of 2D longitudinal image planes from a representative 3D virtual needle biopsy (i.e. needle biopsy IV) is illustrated in Figure 8. The same staging process and the CPA quantification analysis are applied to the resulting 2D longitudinal images. The highest staging score of all 2D image planes is used as the diagnostic staging score. The resulting overall MAD computed with these 2D longitudinal image planes is 0.187 and 1.066 by the Scheuer and Ishak staging method, respectively. With Scheuer staging scores, 37 out of 198 2D longitudinal images deviate from the diagnostic staging score, 1 out of 18 needle biopsies is not consistent with the diagnostic score, and 4 out of 18 3D virtual needle biopsies present internal 2D image planes with diverse staging scores. By the Ishak staging method, 171 out of 198 2D images deviate from the diagnostic staging score, 14 out of 18 needle biopsies are not consistent with the diagnostic score, and 6 out of 18 3D virtual needle biopsies present internal 2D longitudinal image planes with distinct staging scores.

**Figure 8:**
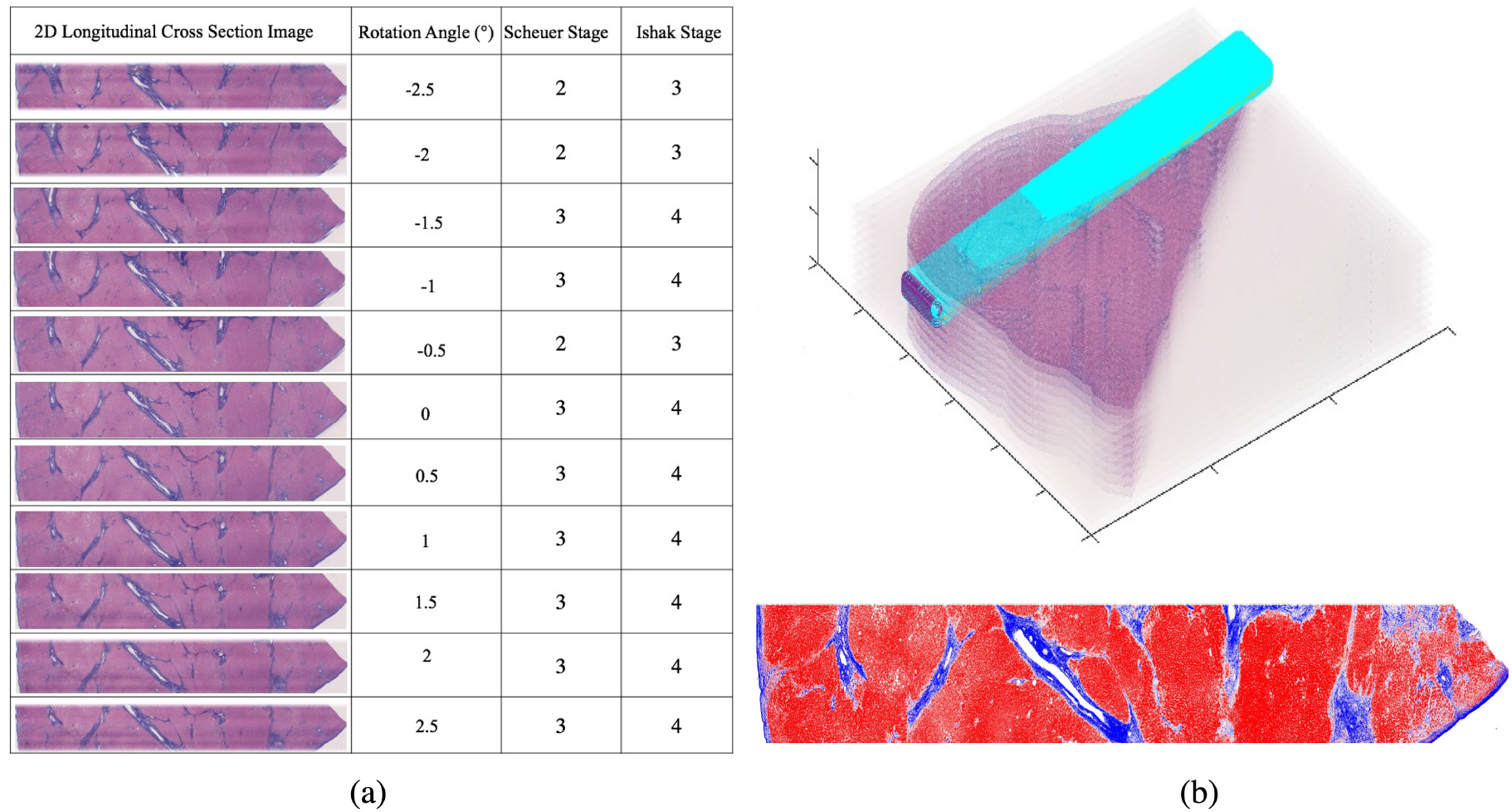
Liver staging difference within a representative virtual 3D needle biopsy. (a) 2D longitudinal cross-sections from a representative 3D virtual needle biopsy (i.e. biopsy IV) with the rotation angle 0°, ±0.5°, ±1°, ±1.5°, ±2°, and ±2.5° are illustrated with their staging scores given by a pathologist; (b) The sampling location and angle of the resulting 3D virtual needle biopsy is demonstrated, along with the labeled 2D longitudinal image with collagen in blue and other tissue components in red, respectively.

To further characterize inter-needle biopsy variation, we next represent each 3D virtual needle biopsy by its 2D image planes rotated around the longitudinal needle axis. Analysis of Variance (ANOVA) is used to test if the average staging scores and CPAs from such 2D longitudinal images are equal across diverse biopsies. The resulting ANOVA test suggests that not all biopsy group means are equal by either staging scores or CPAs. This indicates the presence of tissue heterogeneity in 2D longitudinally rotated biopsy images grouped by virtual needle biopsies. Tukey’s multiple comparison test is next used to identify significantly different biopsy pairs at the significance level 0.05. The total number of different biopsy pairs can be computed by the combinatorics (i.e. combination of 18 taken 2). Demonstrated in Figure 9, 101 out of 153 biopsy pairs (i.e. 66%) are significantly different by the CPA average. By the Scheuer and Ishak staging scores, 109 (i.e. 71%) and 79 (i.e. 52%) out of 153 biopsy pairs present significantly different population means. All these experiments manifest the large collagen heterogeneity captured by 2D longitudinal biopsy images from distinct 3D virtual needle biopsies.

**Figure 9:**
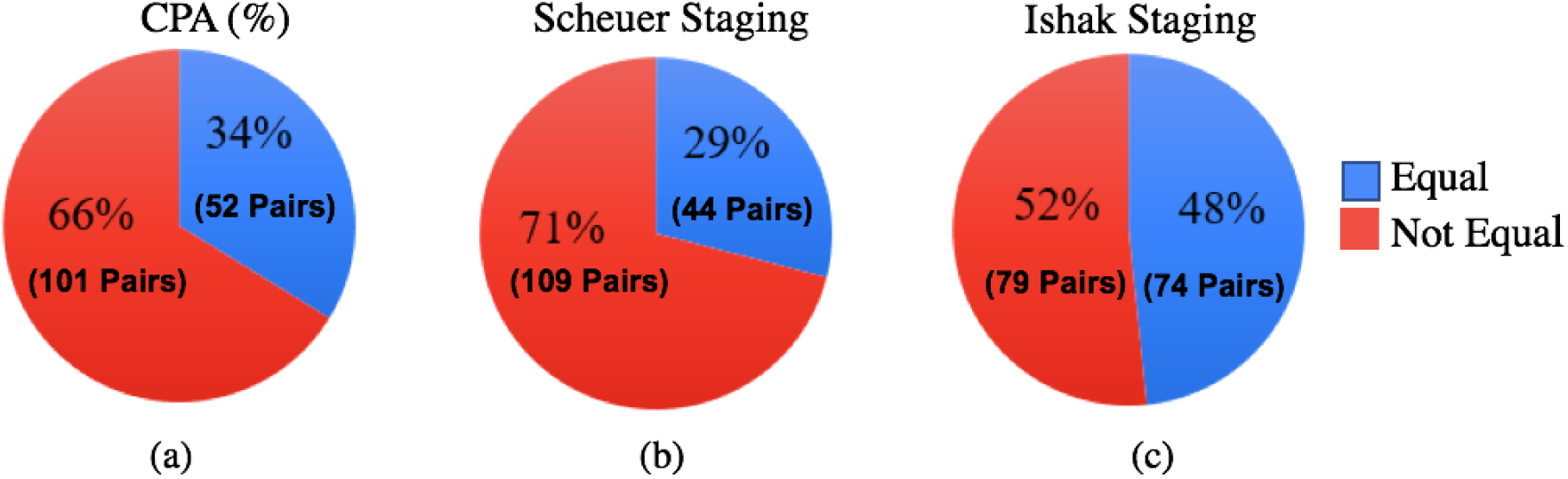
Inter-needle biopsy heterogeneity using 2D longitudinal biopsy images. We represent each 3D virtual needle biopsy by its 2D image planes rotated around the longitudinal needle axis. Analysis of Variance (ANOVA) test suggests that not all biopsy group means are equal by either staging scores or CPAs. Tukey’s multiple comparison test is used to identify significantly different biopsy pairs at the significance level 0.05. The total number of different biopsy pairs can be computed by the combinatorics (i.e. combination of 18 taken 2). 101 out of 153 biopsy pairs (i.e. 66%) are significantly different by the CPA average. By the Scheuer and Ishak staging scores, 109 (i.e. 71%) and 79 (i.e. 52%) out of 153 biopsy pairs present significantly different population means.

### 3.5 Intra-3D virtual needle biopsy sampling heterogeneity

#### Intra-3D virtual needle biopsy heterogeneity by staging scores

The maximum score difference across 2D longitudinal image planes from each 3D virtual needle biopsy is calculated by both Scheuer and Ishak staging methods. We notice a clear staging score difference across internal 2D longitudinally rotated images. There are 4 and 6 out of 18 3D virtual needle biopsies with inconsistent scores from their internal 2D longitudinal image planes by Scheuer and Ishak staging methods, respectively. Taking the highest of the staging scores of all internal 2D images within each 3D virtual needle biopsy, we have one, three, ten, and four 3D virtual needle biopsies with Scheuer-Ishak score pair (2, 4), (3, 3), (3, 4), and (3, 5), respectively. These results suggest a substantial sampling variation within each sampled 3D virtual needle biopsies.

#### Intra-virtual needle biopsy heterogeneity by CPAs

Furthermore, we have studied the intrabiopsy liver CPA heterogeneity with reference to the CPA from the entire 3D liver tissue volume. For each virtual needle biopsy, we test if the average CPA from 2D longitudinal images within a 3D virtual needle biopsy is equal to the overall CPA from the 3D tissue volume. With t-test, we find 14 out of 18 needle biopsies present a significant difference by CPA. This analysis suggests the CPAs from 2D longitudinally rotated images of individual 3D virtual needle biopsies are significantly different from the overall 3D liver tissue volume CPA in most cases. We also use the t-test to determine if the average CPA from 2D longitudinal images of a 3D virtual needle biopsy can represent that of the corresponding 3D virtual needle biopsy. Such analysis results in a p-values less than 0.05 for 17 out of 18 3D virtual needle biopsies. Therefore, there is a significant difference between the CPA derived from individual 3D needle biopsy and that from its 2D longitudinal images. Such analyses suggest that resulting 2D longitudinal images are not good surrogates for the sampled 3D virtual needle biopsies for collagen quantification, unveiling a substantial intra-biopsy liver CPA heterogeneity.

#### 3.6 CPAs of 2D longitudinal image planes across staging groups

We present and compare CPAs of 2D longitudinal image planes across staging groups by two staging methods in Figure 10. By Scheuer staging method, the average biopsy CPAs from biopsies of staging score 2 and 3 are 12.75% and 14.29% respectively with p-value of 4.2e*−*3 by t-test. By contrast, the average biopsy CPAs from biopsies of Ishak stage score 2, 3, 4 and 5 are 17.03%, 13.26%, 14.09% and 13.35% respectively with p-values of 0.254 by ANOVA. Overall, CPA measures are positively correlated with fibrosis stages by Scheuer but not Ishak method.

**Figure 10:**
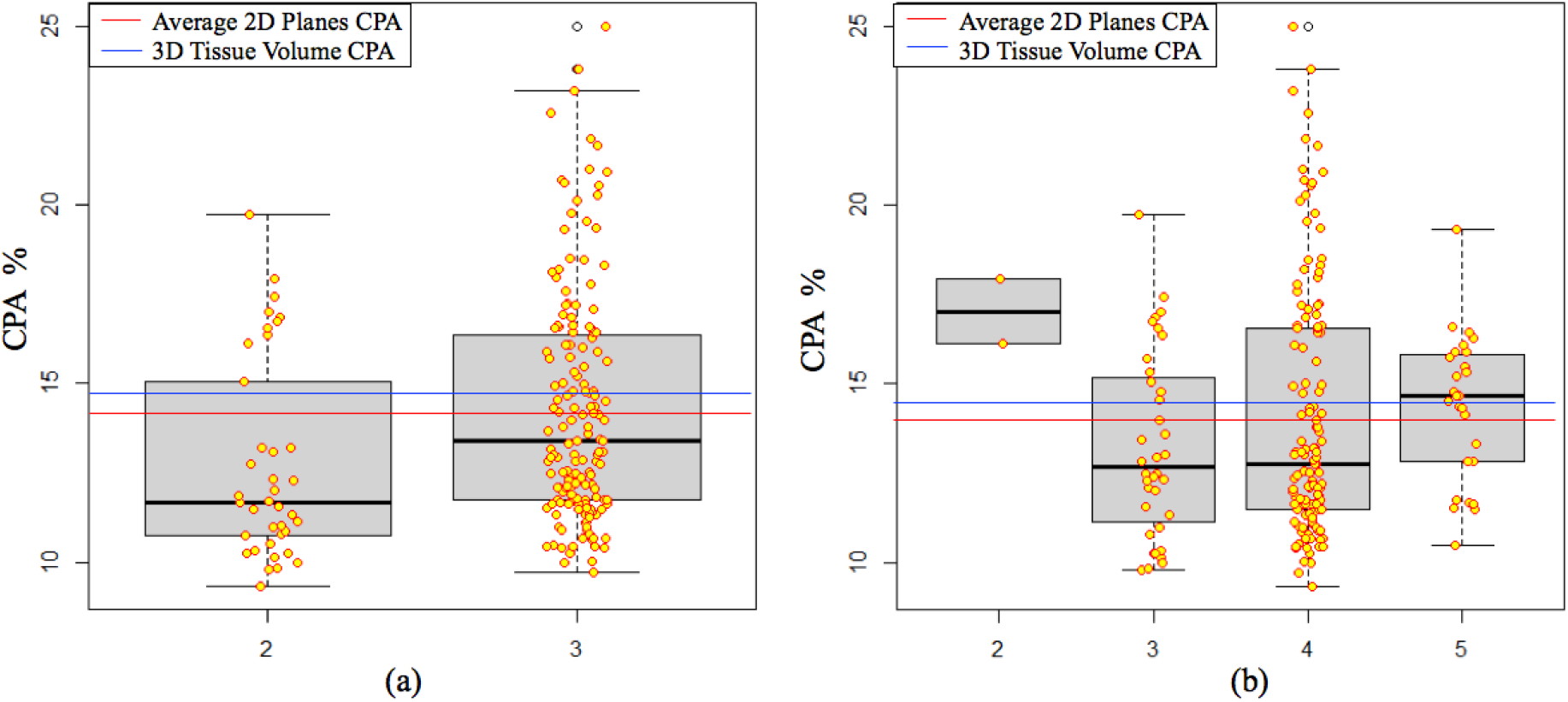
CPAs of 2D longitudinal image planes in 3D virtual needle biopsies across staging groups. Liver CPA quantitation results and their distributions in Scheuer (a) and Ishak (b) staging groups are presented along with the average biopsy CPA (red line) and the overall 3D tissue volume CPA (blue line). By Scheuer staging method, the average biopsy CPAs from biopsies of staging score 2 and 3 present a p-value of 4.2e-3 by the t-test, while those by Ishak stage scores present a p-values of 0.254 by ANOVA.

## 4 Discussion

The liver needle biopsy review is the current gold standard for liver fibrosis and live disease diagnosis. Thus, it is important to have a systematic method to investigate such needle biopsy sampling variation and assess its impact on the resulting diagnostic results. However, there is a paucity of methods for sampling variation studies as it is not practical to take a large number of needle biopsies from the same patient for such investigations. Compared to the limited biopsy samples for diagnosis, larger volumes of tissues resected during surgery operations are more appropriate to support biopsy sampling studies. In this work, we present a new computational analysis pipeline that enables an automatic needle biopsy sampling investigation with highresolution WSIs of serial tissue sections. Multiple virtual 3D needle biopsies are digitally sampled from the 3D liver tissue volume recovered by spatially registered serial WSIs. In each 3D virtual needle biopsy, CPA, a collagen measure strongly correlated with the liver fibrosis stage, is automatically computed by the Masson’s trichrome signal. We present sampling heterogeneity by comparing staging scores and CPAs from random 3D virtual needle biopsy samples and their internal 2D longitudinal images. Experiment results suggest that our method is effective for liver needle biopsy sampling study and can automatically quantify liver collagen components. The developed virtual needle biopsy sampling method and liver collagen quantification approach can be applied to 3D tissue image volumes repeatedly. Therefore, they can support an efficient, reliable and accurate biopsy sampling bias analysis. With minor changes, this analysis method can be extended to a large spectrum of other tissue related investigations where the sampling variation in 3D tissue histology components can significantly affect the clinical reviewing results.

The suite of analysis tools, including serial WSI registration method, 3D virtual needle biopsy sampling function, internal needle biopsy cross-section image generation, and needle biopsy sampling error investigation, are generic and can be further upgraded. Although we specifically demonstrate our needle biopsy sampling investigation on liver tissues, the proposed analysis methodology can be enhanced and customized to a large number of other tissue-related research investigations and clinical applications. Our study represents a proof of concept for the technical challenging problem on 3D digital histology tissue analysis and enables definitive investigations on needle biopsy sampling heterogeneity in a computational manner.

## Demonstration website

https://dplab.gsu.edu/demo.

## Conflict of Interest Statement

The authors have declared no conflicts of interest.

## Author contributions

Y.J. and J.K. conceived the original idea; Y.C. and H.C. acquired and selected image data. Y.J. and J.K. designed the research; Q.L., F.W., S.W., A.B.F. Y.J. and J.K. performed the research; All authors were involved in planning for the project; J.K., Y.J. and Q.L. worked on the manuscript with support from F.W., S.W. and A.B.F. All authors reviewed the final manuscript.

---

**Figure S1:**
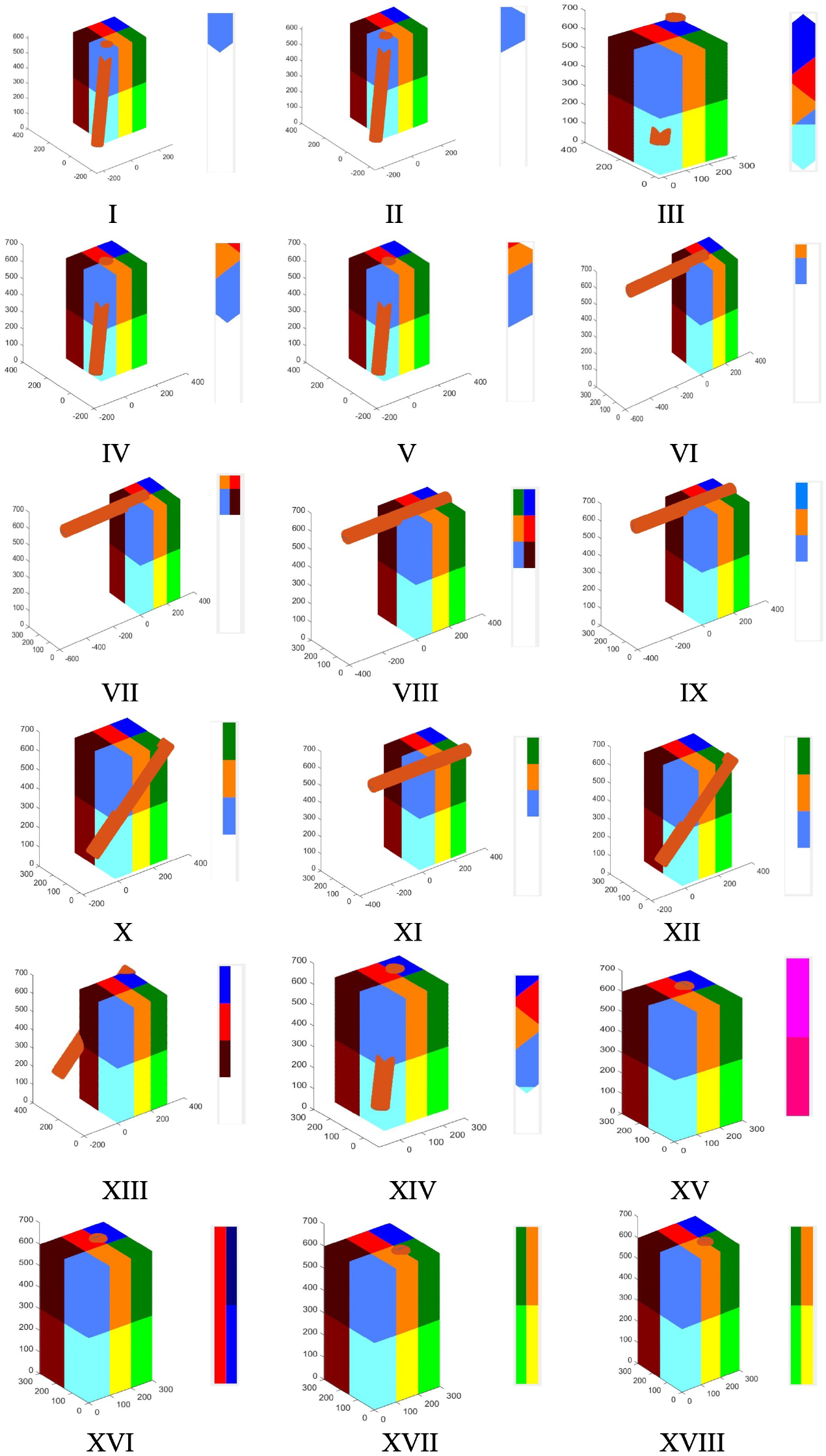
Validation of virtual 3D needle biopsy sampling function with synthetic 3D data volumes. An array of virtual 3D needle biopsy samples over a synthetic data volume of size 300×300×600 voxels is presented. For sampling function validation, each 3D data volume includes 12 cuboids in distinct colors. The resulting 2D longitudinal planes from the corresponding virtual 3D needle biopsies are illustrated next to each 3D synthetic data volume.

**Figure S2:**
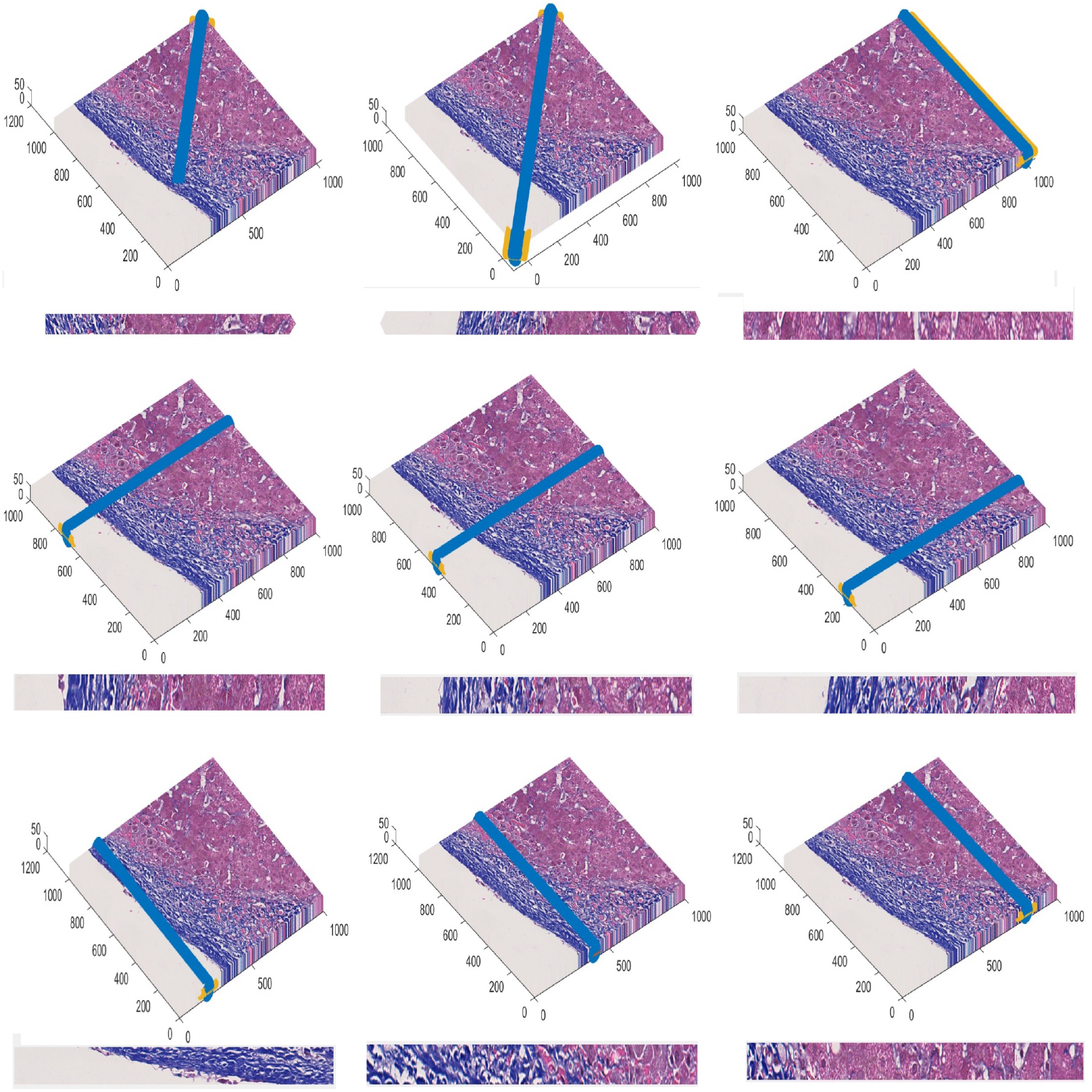
Validation of virtual 3D needle biopsy sampling function with semi-synthetic 3D liver tissue volumes. We construct a semi-synthetic data volume by replicating a tissue region (1, 000 *×* 1, 000) from a WSI for 50 times. Different virtual 3D needles are digitally injected into the semi-synthetic tissue volume, and their 2D longitudinal image views are illustrated and visually validated.

**Figure S3:**
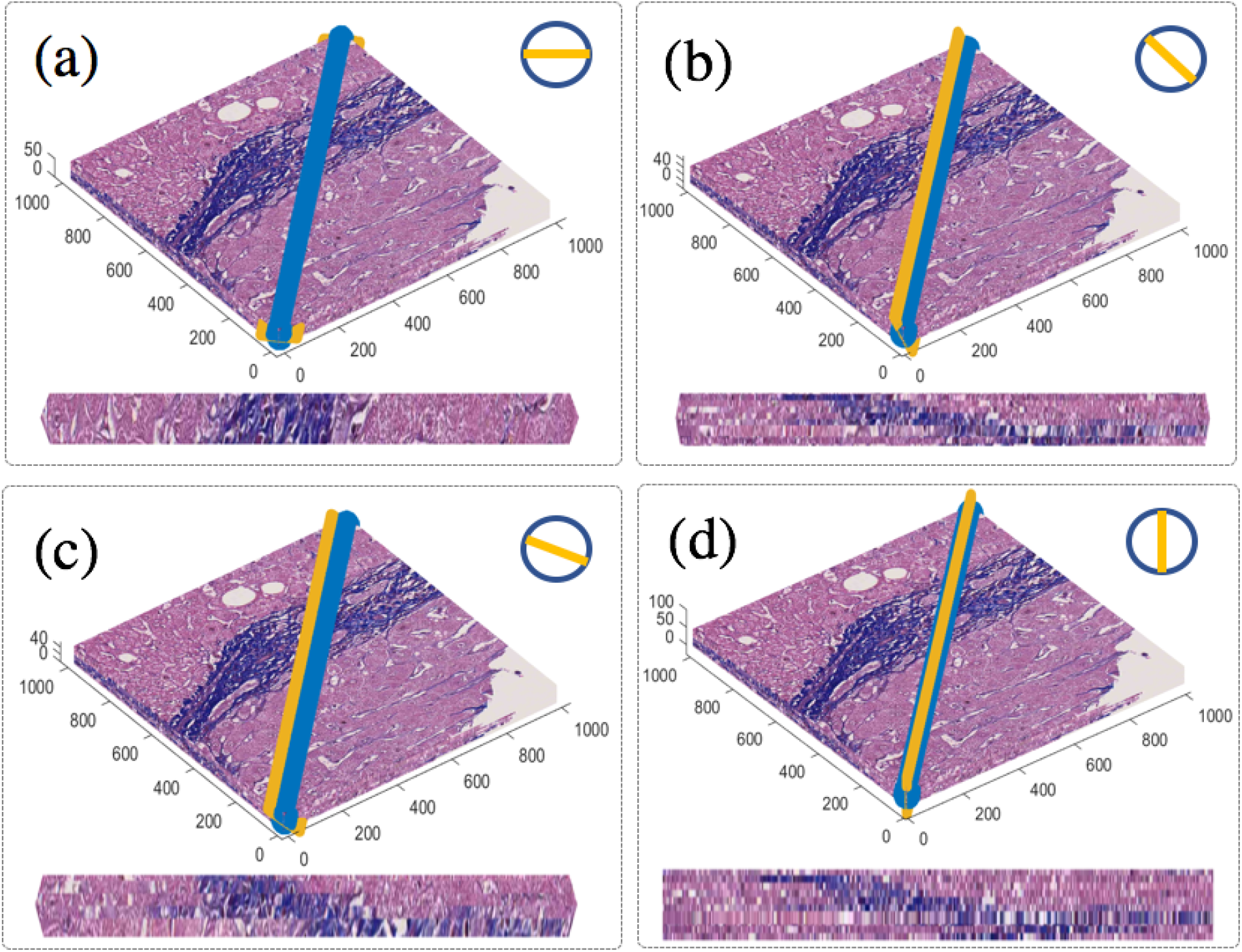
Validation of 2D longitudinally rotated image generation with semisynthetic liver tissue volumes. We present interpolated 2D image planes rotated around the 3D virtual needle axis by the rotation angle (a) 0°, (b) 30°, (c) 60°, and (d) 90°, respectively. The corresponding 3D virtual needles in reference to the 3D semi-synthetic tissue volumes are also presented, with virtual needle biopsy sampling parameters (*P*_0_ = [500, 100, 25]; *L* = 1, 500; *R* = 25; *θ* = 90^°^; *φ* = 45^°^).

**Figure S4:**
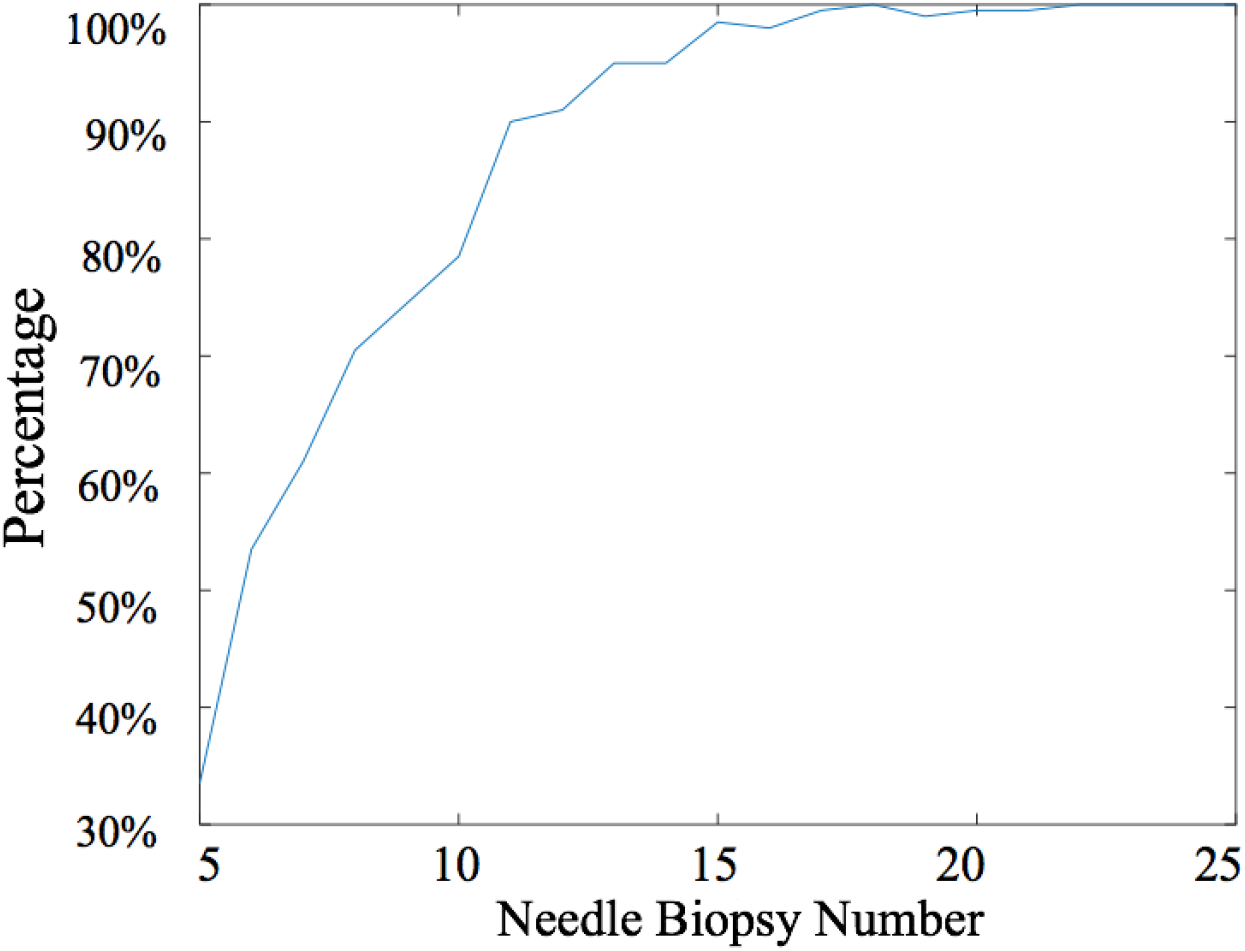
Selection of virtual 3D needle biopsy number for sampling heterogeneity investigation. To determine the optimal biopsy number, we vary the number of virtual needle biopsies in sampling simulation tests. For each fixed biopsy number, we repeat the virtual sampling process at randomly selected needle locations and angles for 200 times. After each set of 200 virtual sampling processes, the resulting CPAs from individual virtual needle biopsies are compared with the overall liver CPA from the reconstructed virtual 3D liver tissue volume by t-test. We use the percentage of the sampling processes with the p-value ¡ 0.05 as a measure of tissue heterogeneity representation power. Such a measure is plotted as a function of the needle biopsy number.

## Notes

### Competing Interest Statement

The authors have declared no competing interest.

